# Large gradient of susceptibility to esca disease revealed by long-term monitoring of 46 grapevine cultivars in a common garden vineyard

**DOI:** 10.1101/2024.03.11.584376

**Authors:** Pierre Gastou, Agnès Destrac Irvine, Clarisse Arcens, Eva Courchinoux, Patrice This, Cornelis van Leeuwen, Chloé E. L. Delmas

## Abstract

Grapevine (*Vitis vinifera* L.) is prone to many fungal diseases, including esca, a severe vascular disease threatening the wine sector and for which there is no cost-effective cure. Susceptibility to esca varies between cultivars in different infection conditions. It may therefore be possible to use the genetic diversity of grapevine cultivars to mitigate disease impact. However, the genetic component of esca susceptibility has rarely been investigated in the vineyard, and the specific mechanisms and varietal traits underlying esca susceptibility remain unknown.

In this study, we monitored the incidence and severity of esca foliar symptoms and plant dieback (apoplexy and mortality) at plant level for seven years, on 46 cultivars planted in an experimental common garden, to separate the genetic component of esca susceptibility from the effects of environment and cropping practices. We observed a broad gradient of varietal susceptibility, with a mean incidence of 0 to 26% of vines expressing esca foliar symptoms depending on the variety. This gradient remained similar across years and, unlike the severity of foliar symptoms, the incidence of grapevine dieback was significantly correlated with that of foliar symptoms. We detected a significant but weak and very localised phylogenetic signal for the incidence of esca foliar symptoms in this panel of cultivars.

We then explored the relationships between epidemiological metrics and ecophysiological and phenological traits phenotyped on the same plot. Esca disease incidence was negatively correlated with δ13C across cultivars, suggesting that varieties with higher water use efficiency are less prone to the expression of esca symptoms on leaves. Moreover, the least vigorous cultivars were among the least susceptible, although this relationship was not significant. By contrast, neither phenological stages nor nitrogen status were significantly predictive of cultivar susceptibility to the disease.

Together, these results provide new insight into the potential of genetic resources for use in the sustainable management of grapevine trunk diseases and open up new perspectives for studying the pathological and physiological determinants of their incidence.

## Introduction

The domesticated grapevine *Vitis vinifera* L. ssp. *vinifera* has a high level of genetic and phenotypic diversity, generated naturally by recombination, hybridisation, and mutation events, as well as through human-assisted selection. More than 5,000 cultivars are currently registered (This *et al*., 2006), although only a small proportion are widely cultivated worldwide (Anderson and Aryal, 2013). Cultivar diversity is now considered an excellent tool for managing abiotic and biotic pressures in a sustainable manner (Merdinoglu *et al*., 2018; Wolkovich *et al*., 2018). There is therefore a need to improve our understanding of the variability between cultivars by phenotyping them for a wide range of traits. Several stress-related traits, including carbon isotope discrimination in berry juice at maturity (δ13C; Plantevin *et al*., 2022), cold hardiness (Ferguson *et al*., 2014), and susceptibility to pests and diseases (Boso *et al*., 2011; Gaforio *et al*., 2011; Paňitrur-De La Fuente *et al*., 2018), have already been studied in a range of cultivars. The susceptibility of the grapevine vascular system to various stresses has also recently been studied. Xylem and hydraulic traits have been phenotyped in a range of *Vitis* genotypes of different origins and levels of drought tolerance (Dayer *et al*., 2022; Lamarque *et al*., 2023) or resistance to vascular pathogens (Pouzoulet *et al*., 2020; Fanton and Brodersen, 2021).

Vascular biotic stresses include grapevine trunk diseases, which are responsible for substantial yield losses in vineyards worldwide (Gramaje *et al*., 2018). One such disease is esca, which is detected in the vineyards through the summer expression of foliar symptoms and yield losses (Lecomte *et al*., 2012). Esca pathogenesis has been associated with various types of trunk necrosis (Mugnai *et al*., 1999) probably involving a complex community of fungal pathogens (Bruez *et al*., 2014). This community includes a few key pathogens from both Ascomycota (including *Phaeomoniella chlamydospora* and *Phaeoacremonium minimum*) and Basidiomycota (e.g. *Fomitiporia mediterranea*). However, no direct role in foliar symptom onset has ever been demonstrated for any of these pathogens, which are present in both healthy and necrotic wood tissues (Bruez *et al*., 2016) and in both asymptomatic and symptomatic plants (Hofsteter *et al*., 2012; Bruez *et al*., 2014). These fungi probably also interact with bacteria (Bruez *et al*., 2020) and seem to be restricted to the perennial organs (Bortolami *et al*., 2019).

Foliar symptoms are associated with xylem hydraulic failure and impaired photosynthesis (Bortolami *et al*., 2019, 2021a; Ouadi *et al*., 2021; Dell’Acqua *et al*., 2023). The incidence of esca foliar symptoms is probably affected by multiple factors (as reviewed in Claverie *et al*., 2020), including plant age, pedoclimatic conditions, viticulture practices, the plant- and soil-associated microbiota, and the plant material. The genetic diversity of the grapevine response to esca may be associated with differences in phenotypic characteristics (ecophysiology, phenology) at cultivar level. Based on what we now know about plant-to-plant variability within a single cultivar, mineral status, water status and plant phenology are promising avenues to be explored. The incidence of esca increases following foliar applications of nutrient sprays, which suggests that a higher nutrient content in the leaves may be associated with a higher incidence of esca (Calzarano *et al*., 2009). High levels of grapevine transpiration may facilitate the translocation of pathogenic toxins and metabolites throughout the vine (Bortolami *et al*., 2021b). Differences in phenology can interfere with the co-ordination between ontogenic susceptibility and favourable climatic periods for disease development (Serra *et al*., 2018).

The susceptibility of grapevine cultivars to trunk pathogens and diseases, especially esca, has been assessed with three methods. The first one reports varietal susceptibility to fungal infection and involves phenotyping the internal necrotic lesions that develop after artificial inoculation with one or more fungal species, such as *P. chlamydospora* (e.g. Pouzoulet *et al*., 2017, 2020; Martínez-Diz *et al*., 2019) and, to a lesser extent, *P. minimum* (Feliciano *et al*., 2004; Gubler *et al*., 2004). However, the results obtained by this method are not readily transferable to field conditions, particularly as fungal inoculations do not reproduce foliar symptoms (Claverie *et al*., 2020). The second method is the monitoring of esca incidence (e.g. proportion of plants presenting foliar symptoms or dieback) in a network of productive vineyards, over an entire region, or even at the national scale (Bruez *et al*., 2013). However, one of the major limitations of this approach is the large number of confounding factors, such as viticulture practices, climatic conditions, soil type, plant age, and rootstock. These biases can be overcome, making it possible to compare grapevine cultivars within the same climatic and cultural context, by monitoring a single experimental vineyard planted with a set of cultivars over multiple years. In the last decade, such experimental setups have been used to compare numerous cultivars in common environments (e.g. Murolo and Romanazzi, 2014). However, no experiment to date has been purposely designed to take into account pedoclimatic microvariability and the origin of the plant material, both of which could account for a significant amount of the variability in disease susceptibility (Kovács *et al*., 2017; Gramaje *et al*., 2018). As a result, limited information is currently available regarding cultivar differences in terms of the incidence of esca foliar symptoms and dieback in a common environment. Are the differences between cultivars consistent between years? What are the ecophysiological drivers of esca susceptibility at cultivar level?

In this study, we focused on the genetic component of grapevine susceptibility to esca foliar symptoms and dieback, independent of *terroir* effects, and on the mechanisms potentially underlying differences between cultivars. We monitored the incidence and severity of esca foliar symptoms and plant dieback, and a range of phenological and ecophysiological traits in a common garden experimental vineyard planted with 46 cultivars, over periods of seven and six years, respectively. This design made it possible to study cultivar-specific variability without bias due to other factors, such as year, soil, plant material origin, and viticulture practices. We first investigated the range of cultivar susceptibility to esca foliar symptoms and grapevine dieback. We then explored multiple correlations between different traits and years, to assess the temporal consistency of the ranking of varieties for susceptibility. Finally, we investigated the relationships between esca foliar symptoms and dieback on the one hand, and phenological and ecophysiological traits, such as key phenological stages, nitrogen status at flowering, pruning weight, and δ13C, a proxy of water use efficiency, on the other.

## Materials and methods

### 1. Common garden experimental vineyard

The VitAdapt vineyard (as described by Destrac-Irvine and van Leeuwen, 2016) was designed as a common garden, and is located at the *Institut National de Recherche pour l’Agriculture, l’alimentation et l’Environnement* (INRAE) research station (Villenave d’Ornon, Nouvelle-Aquitaine, France), at 44°47’23.83 N’’, 0°34’39.3’ W’. This vineyard, on a sandy-gravel soil, was planted with 52 genotypes (47 *Vitis vinifera* L. cultivars and 5 *Vitis* hybrids). In this study, 45 *V. vinifera* L. cultivars, and one hybrid of several *Vitis* species — Hibernal, an F2 progeny of a Seibel 7053 x Riesling cross (Vitis International Variety Catalogue - VIVC; www.vivc.de) — were monitored. Among them, 28 were red-berried cultivars and 18 were white-berried cultivars (Supplementary Table S1). All plants were grafted onto Selection Oppenheim 4 (SO4) clone 761 rootstock and planted at a density of 5,555 vines/ha, corresponding to a spacing of 1.8 m between adjacent rows and 1 m between adjacent vines. Eight cultivars were first planted in 2010, all the others being first planted in 2009 (Supplementary Table S1). Each cultivar was tested for major viral diseases before planting, and only non-contaminated material was planted. All vines were pruned according to the double guyot system and grown without irrigation. Pests and diseases were controlled by an integrated management programme, and weeds were controlled by mechanical tillage beneath the row. No specific management of trunk diseases was carried out. Cover crops were planted in every other row, alternating from year to year. The VitAdapt vineyard was organized into a five-randomised block design to account for variability in soil properties and pathogen inoculum. Thus, we assume that all cultivars were exposed to similar risks of pathogen infection. Each block comprised one subplot per cultivar, in which 10 vines were planted in two adjacent rows. In this study, only four blocks were monitored, resulting in a total of 184 subplots (1,840 vines; 40 per cultivar, 46 cultivars) subjected to ecophysiological monitoring and esca and dieback monitoring for six and seven years, respectively. No artificial inoculation of trunk pathogens has been conducted in this vineyard, and dead vines were regularly removed to prevent local hotspots of spore production.

### 2. Monitoring of foliar symptoms of esca and plant dieback

We monitored 1,840 vines individually by eye, to check for esca foliar symptoms (examples are shown in Figure 1) and plant dieback (apoplexy and death), between 2017 and 2023. Evaluations were performed one to three times per year, between early July and mid-September. For each monitoring campaign, all the plants were inspected during a period of no more than one week. For each vine, a score was assigned at individual arm level for the observed phenotype (asymptomatic, presence of esca foliar symptoms/symptomatic, apoplectic, dead, young replanted vine, missing). When foliar symptoms were observed, severity was scored by determining, for each arm, the proportion of symptomatic shoots and the degree of leaf and berry dehydration (as described in Supplementary Table S2). Briefly, the score attributed for each arm was either1 (light symptoms), 3 (moderate symptoms) or 5 (severe symptoms, i.e. ‘tiger-striped’ leaves and dried berries). In years with multiple assessments, the maximum severity index was retained. To assign a score for each vine, we averaged the score for each of its symptomatic arms.

**Figure 1.**
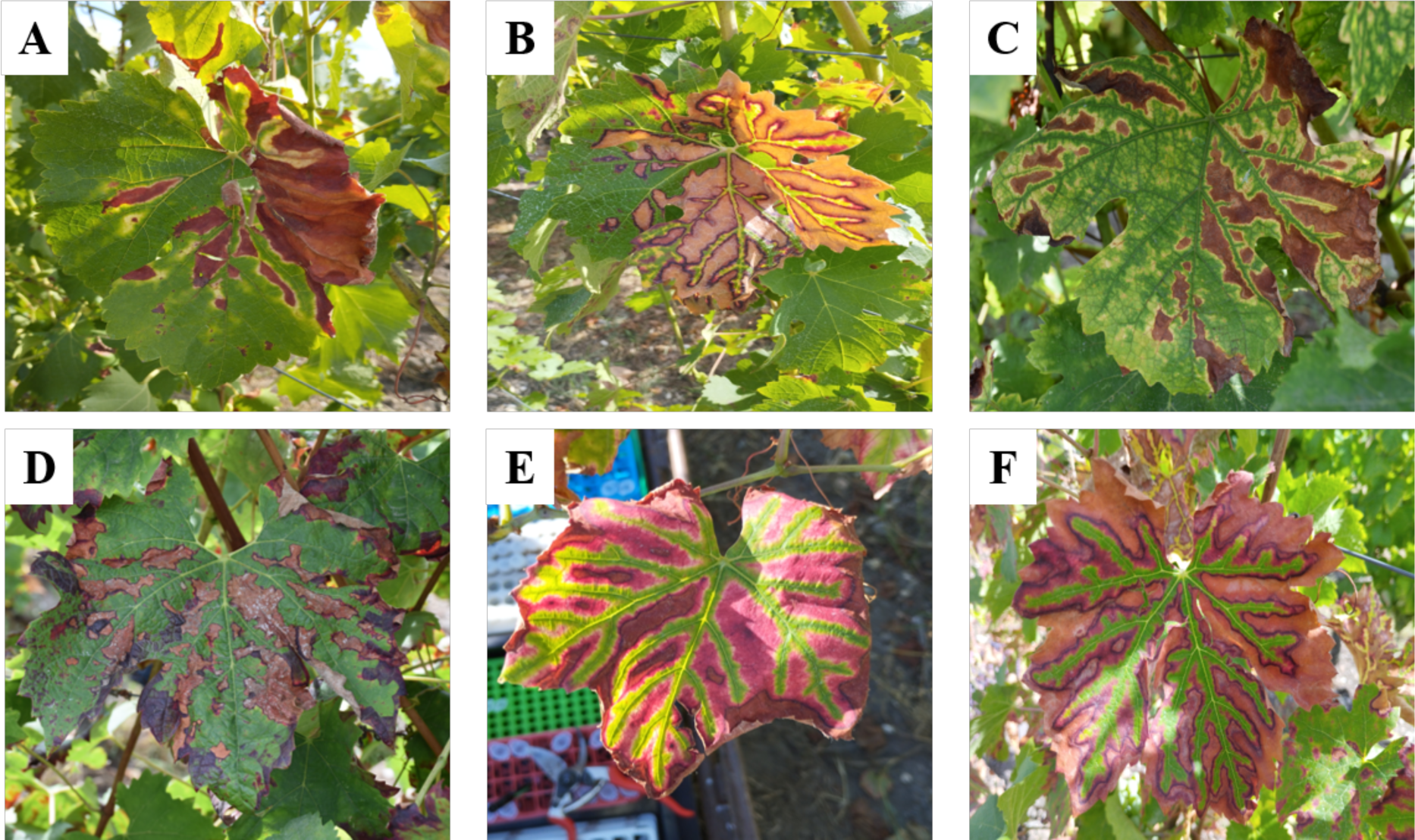
Examples of various foliar phenotypes for esca on different *Vitis vinifera* cultivars grown in a common garden experimental vineyard. A-C: white-berried cultivars (A) Muscadelle; (B) Chasselas; (C) Viognier. D-F: red-berried cultivars (D) Alvarinho; (E) Pinot Noir; (F) Cabernet Franc. Note that the disease status of a vine is determined by phenotyping the entire shoot, not just individual leaves.

Three epidemiological metrics were calculated from the data collected at subplot scale for esca foliar expression and vine dieback. The incidence of esca foliar symptoms was calculated as the proportion of the total number of productive plants (i.e. excluding totally apoplectic and dead plants, as well as young replanted vines) displaying typical esca foliar symptoms. The cumulative incidence of foliar symptoms was also calculated as the proportion of the productive vines having expressed esca foliar symptoms in at least one of the seven years monitored. The mean severity of esca foliar symptoms was calculated at subplot scale, by averaging the score attributed to each symptomatic vine of the subplot. Finally, the incidence of plant dieback was calculated as the proportion of mature plants (i.e. excluding young replanted vines) with at least one unproductive arm (i.e. apoplectic or dead). During the plant phenotyping process in the field, no *a priori* assumption was made concerning the cause of plant dieback, which may have multiple abiotic or biotic causes.

### 3. Phenotyping of ecophysiological and phenological traits

We explored the mechanisms underlying cultivar susceptibility to esca and dieback, using cultivar phenology data and monitoring data for three ecophysiological traits collected at the subplot scale in the VitAdapt vineyard.

#### 3.1. Monitoring of phenological stages

The phenological evaluation was performed visually, in the field, on 10 vines of each cultivar in each of the four blocks, as described by Destrac-Irvine *et al*. (2019). Three different stages were monitored: bud break, flowering and veraison. Notations were based on the BBCH scale (Lorenz *et al*., 1995). The bud-break stage corresponds to the date on which 50% of the buds have reached the BBCH 07 stage; the flowering stage corresponds to the date on which 50% of the flowers have reached the BBCH 65 stage; and the veraison stage corresponds to the date on which 50% of the berries have reached the BBCH 85 stage. Six years of data were included in this study, from 2017 to 2022.

#### 3.2. Carbon isotope discrimination (δ^13^C)

We determined δ^13^C levels in berry juice sugars obtained each year, at maturity, from 10 vines of each cultivar in each of the four blocks, as described by Plantevin *et al*. (2022). All samples were analysed in an external laboratory (UMR CNRS/Plateforme GISMO UMR 6282 BIOGEOSCIENCES *Université de Bourgogne*, 21000 Dijon, France). Briefly, the juice was extracted and analysed on a Vario Micro Cube elemental analyser coupled in continuous flow mode to an isotopic ratio mass spectrometer (IsoPrime, Elementar). Results are expressed according to the Vienna Pee Dee Belemnite (VPDB) international reference. Six years of data were included in this study, from 2017 to 2022.

#### 3.3. N-tester measurements

N-tester measurements were performed at flowering (as defined above), which is considered to be a key stage for nitrogen assimilation by grapevines (Celette and Gary, 2013), on 30 leaves of each cultivar in each of the four blocks, according to the manufacturer’s protocol (Yara, Oslo, Østland, Norway). Briefly, each leaf was clamped with the device, for the measurement of transmittance (at wavelengths of 650 and 960 nm), which provides a standardised index of chlorophyll content that can be used as a proxy for vine nitrogen status. As a means of accounting for cultivar-specific nitrogen-status behaviour under similar conditions, raw values were transformed according to varietal rounded corrections. The mean value for Sauvignon Blanc was similar to the mean value across all varieties, so this cultivar was used as the reference variety for this index. For each cultivar, the difference between the mean value for all available years for the cultivar (2015-2021 for 35 cultivars; 2020-2021 for the other 11), and the mean value for Sauvignon Blanc was calculated. An Ascendant Hierarchical Classification (AHC) method was then applied to the N-tester values to group the cultivars and obtain an average correction for all the cultivars of a single group. The N-tester value of each cultivar was then corrected with the correction coefficient for its group, as described in Supplementary Table S3. Five years of data were included in subsequent analyses, from 2017 to 2021.

#### 3.4. Pruning weight quantification

Pruning weight was quantified during the winter for all varieties in each of the four blocks. For each cultivar, two measurements were made for each block, by averaging the value three central vines of a row (i.e. six vines per subplot). Only the wood from the previous growing season removed from the plants during pruning was weighed. All measurements were performed during the same period of the year in each year: between January 15th and 30th. Six years of data were included in this study, from 2017 to 2022.

### 4. Data analysis

We first evaluated the effect of block on the three epidemiological metrics: esca foliar symptom incidence, esca foliar symptom severity, dieback incidence. Block was found to have a significant impact on the incidences of both foliar symptoms and dieback (*p* = 0.03 and *p* = 10^−5^, respectively; Supplementary Figure S1). Thus, block was included in subsequent models as a random effect to account for this variability.

We assessed the effects of cultivar and year (entered as fixed effects) on the three epidemiological metrics using three independent mixed modelling procedures, with block as a random effect. The effect of berry skin colour (white/red) on the three epidemiological metrics was independently modelled as a fixed factor, with cultivar and block as random effects.

We investigated the relationships between epidemiological metrics using mean values per cultivar over blocks and years. This made it possible to model the average inter-variety variation of traits independently of temporal and spatial variability. Correlations at cultivar level were assessed in Pearson’s correlation tests (i) between epidemiological metrics, (ii) between epidemiological metrics for each single year, and (iii) between years for each single epidemiological metric. Independent modelling procedures were performed for pairs of variables found to be significantly correlated.

The mean values per cultivar over blocks and years were also used to test the relationships between quantitative ecophysiological and phenological traits on the one hand, and the three epidemiological metrics on the other. Correlations between all pairs consisting of one epidemiological and one phenotypic variable were tested in Pearson’s correlation tests. Independent modelling was performed for all pairs of variables found to be significantly correlated, with the epidemiological metric modelled as the response variable, and the ecophysiological or phenological traits as fixed effects.

For all these analyses, linear models (LM) or linear mixed models (LMM) were used for response variables following a normal distribution (i.e. foliar symptom severity). For binomial response variables (i.e. the incidences of foliar symptoms and dieback), generalised linear models (GLM and GLMM) were fitted, using the binomial family with “cloglog” links (accounting for a non-symmetric distribution). Each model was graphically validated according to the normality of the residuals (QQ-plot) and the homogeneity of the residual variance (residuals vs. fitted, residuals vs. predictors). All analyses involving epidemiological metrics were conducted for the period 2017-2023, whereas analyses including other phenotypic traits were conducted for the period 2017-2022. Analyses were performed with R v.4.2.1 software (R Core Team, 2022) and the RStudio interface. Linear modelling was performed with the “lme4” package (Bates *et al*., 2015), and model validation was performed with “DHARMa” (Hartig and Lohse, 2022). Correlation analyses were performed with the “corrplot” package (Wei, Simko, 2021).

### 5. Phylogenetic signal based on epidemiological metrics

We built a phylogenetic classification of the 46 cultivars included in this study based on grapevine genotyping data for 20 microsatellite markers (SSR), as described by Laucou *et al*. (2011). We used the genetic distances between cultivars obtained from different hierarchical clustering analyses: two classical methods for statistical clustering (i.e. Euclidean and Ward) and two methods adapted for phylogenies (i.e. unweighted pair group method with arithmetic mean (UPGMA) and neighbour joining (NJ)). For phylogenetic signal analysis, we visually inspected the four phylogenies and selected the phylogeny best reflecting the known relationships between cultivars. All information on kinship between the cultivars used in this work was based on the VIVC (www.vivc.de).

We searched for a phylogenetic signal in cultivar susceptibility to esca using methods based on autocorrelations computed both on the global phylogenetic tree and local nodes of the phylogeny, implemented in the “phylosignal” package (Keck *et al*., 2016). A global Moran’s I index was calculated for each epidemiological metric and tested against the null hypothesis of an absence of phylogenetic signal. Phylogenetic correlograms were then constructed to visualise the distribution of this index over a gradient of phylogenetic distances and its confidence envelope based on bootstrapping (1,000 repetitions). The local Moran’s index I_i_ was also plotted alongside the phylogeny to locate autocorrelation patterns more precisely through LIPA analysis. The existence of local phylogenetic signals was assessed in a permutation-based test (999 repetitions).

## Results

### Global incidences of esca and dieback and changes over time

The mean incidence of esca foliar symptoms, for all cultivars, blocks, and years, was 7.3% (standard deviation, SD = 13.6%), with considerable variability (relative standard deviation, RSD = 185.4 %; Table 1). The mean cumulative incidence of esca foliar symptoms (i.e. observed in at least one year) was 30.5%, implying that almost one vine in three expressed esca foliar symptoms in at least one year. This value is five times higher than the mean annual incidence (Table 1). Foliar symptom severity had a mean score of 2.5 and was much less variable than the incidence of foliar symptoms (RSD = 59.4%; Table 1). Plant dieback had a mean incidence of 4.1%, which is lower than that of foliar symptom incidence by a factor of 1.8, but was much more variable (RSD = 233.0%; Table 1). An analysis of the declining vines showed that 16% expressed apoplexy on a single arm, 51% had a single dead arm, and 33% were totally unproductive (i.e. 11% totally apoplectic, 18% totally dead, and 4% with both phenotypes).

**Table 1.** Overall statistics for esca and dieback epidemiological metrics in the VitAdapt common garden experimental vineyard (Villenave d’Ornon, France) between 2017 and 2023. The statistics given are means, standard deviation, relative standard deviation, minimum and maximum. Values are calculated for 1288 observations (46 cultivars; four blocks; seven years). The severity index ranged from 1 (light esca foliar symptom on a single arm) to 5 (severe esca foliar symptom on both arms) and was scored only for plants with symptoms.

| Trait | Statistical individual | mean | SD | RSD | min | max |
| --- | --- | --- | --- | --- | --- | --- |
| Foliar symptom incidence | Observation (n=1288) | 7.3% | 13.6% | 185.4% | 0.0% | 90% |
| Cumulative incidence over seven years | Cultivar (n=46) | 30.5% | 24.0% | 74.4% | 0.0% | 78.5% |
| Foliar symptom severity index | Observation (n=858) | 2.5 | 1.5 | 59.4% | 1.0 | 5.0 |
| Dieback incidence | Observation (n=1288) | 4.1% | 9.6% | 233.0% | 0.0% | 71.4% |

The incidences of esca and dieback increased significantly between 2017 and 2023, with the ageing of the plot (from 7.8 to 13.8 years old on average). This temporal effect was highly significant for the incidence of foliar symptoms (*p* < 10^−16^). The incidences of esca and dieback in 2021 and 2022 were significantly higher than those in the previous four years, with mean values almost six times higher than those obtained in 2017. Finally, the highest incidences occurred in 2023, when the mean incidence of foliar symptoms was more than seven times higher than that in 2017 (Figure 2A). A similar pattern was observed for the incidence of dieback (*p* < 10^−16^). Plant dieback rates were very low during the first four years of monitoring, between 0% and 1%. The proportion of unproductive plants was greatest in 2022 and 2023, when it was significantly higher than in all other years (Figure 2C). The severity of esca foliar symptoms also differed significantly between years (*p* = 10^−11^). However, severity did not follow a specific temporal pattern similar to that for incidence (Figure 2B).

**Figure 2.**
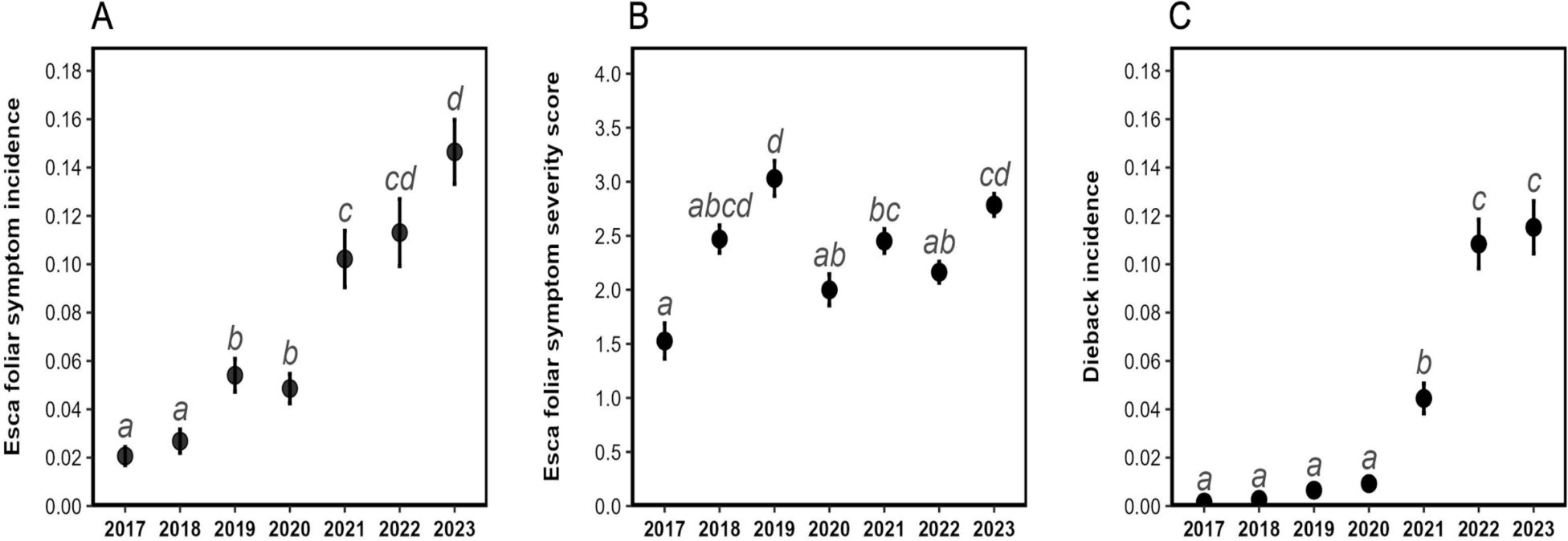
Changes over time in the three variables related to esca foliar symptoms and dieback for all cultivars from 2017 to 2023 (mean plant age increasing from 7.8 years in 2017 to 13.8 years in 2023) (A) Incidence of foliar symptoms of esca (mean ± SEM); (B) Severity of esca foliar symptoms (mean ± SEM), i.e. mean severity index (according to the rating scale described in Supplementary Table S2) calculated on the set of symptomatic plants; (C) Incidence of plant dieback (mean ± SEM). The letters correspond to the groups of significance according to Tukey tests with an alpha risk of 5%.

### Esca incidence differs considerably between cultivars

The incidence of esca foliar symptoms differed significantly between the 46 cultivars monitored from 2017 to 2023 (*p* < 10^−16^). We found a large gradient of susceptibility to esca foliar symptoms, as four grape varieties (Merlot, Petit Manseng, Tannat and Xinomavro) never expressed symptoms during the seven years of monitoring whereas eight cultivars (Cabernet-Sauvignon, Castets, Tempranillo, Sauvignon Blanc, Cabernet Franc, Mourvèdre, Saperavi and Chenin) had mean annual incidences above 20% (Figure 3A).

**Figure 3.**
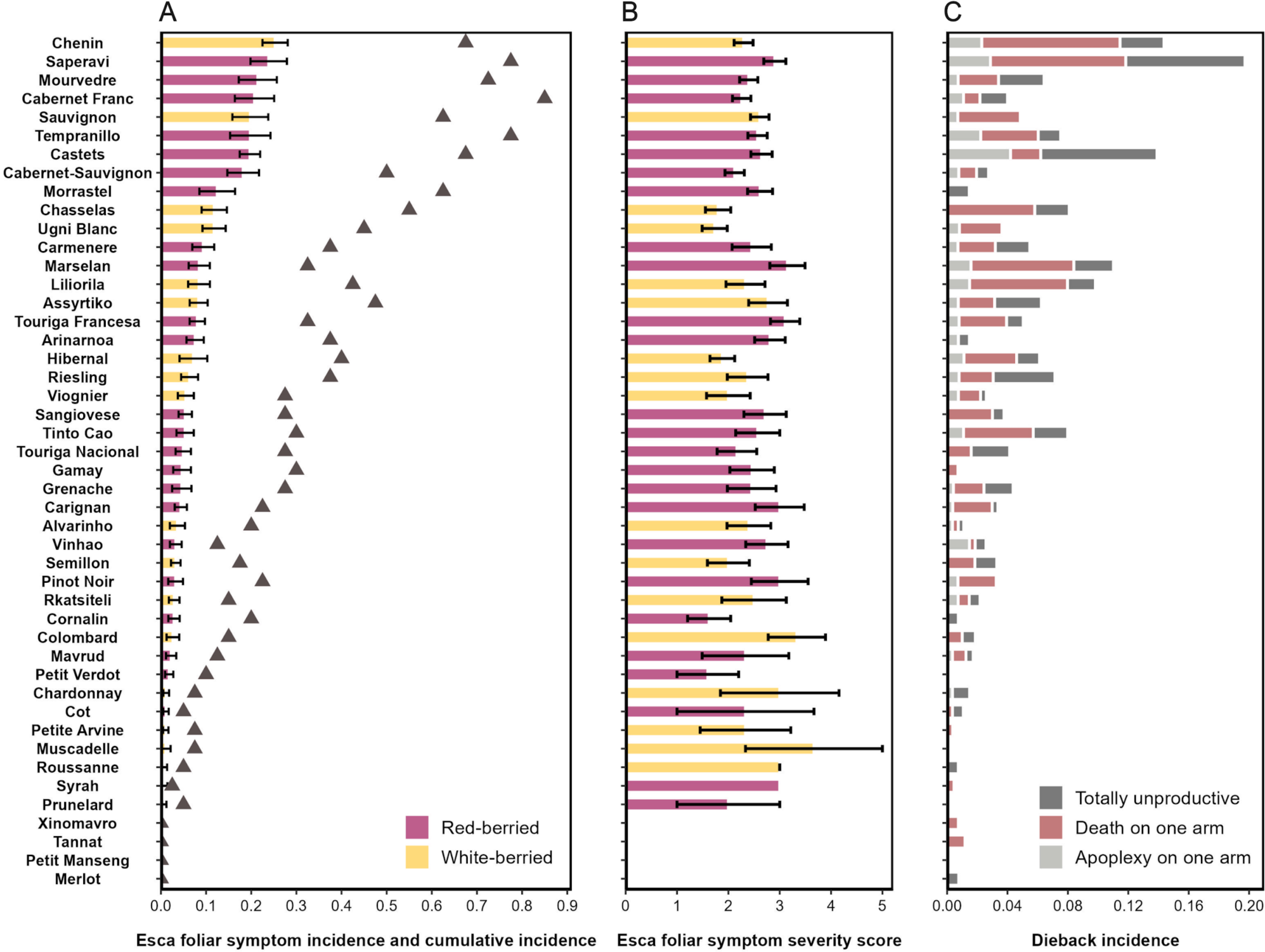
Variability of susceptibility to esca foliar symptoms and dieback among grapevine cultivars for the period 2017 to 2023. (A) Mean incidence of esca foliar symptoms for the 46 cultivars monitored. Bars and error bars represent the means ± SEM and the grey triangles indicate the cumulative incidence over the seven years; (B) Range of severity of esca foliar symptoms in symptomatic plants only (mean ± SEM). The severity index, which ranged from 0 to 5, is described in Supplementary Material 2; (C) Range for the incidence of plant dieback. Light grey corresponds to plants with one apoplectic arm (total dehydration of the canopy). Red corresponds to plants with one dead arm. Dark grey corresponds to totally unproductive plants (apoplexy or death on both arms). Grapevine cultivars are ordered according to the mean incidence of foliar symptoms presented in (A). White-berried cultivars are shown in light yellow and red-berried cultivars are shown in dark purple.

Excluding the four cultivars with an overall incidence of zero, cumulative incidence ranged from 2.5% to 85%, and was 2.7 to 7 times higher than the mean incidence according to the cultivar (Figure 3A). The cumulative incidence of esca was highly correlated with the incidence of foliar symptoms (r = 0.95; *p* < 10^−16^).

The severity of esca foliar symptoms differed significantly between cultivars (*p* = 0.04) but the variability was lower than that for incidence (Figure 3B). Each cultivar presenting esca foliar symptoms could display any degree of disease severity, regardless of its mean incidence of foliar symptoms.

The incidence of plant dieback also differed significantly between cultivars (*p* < 10^−16^). Four cultivars (Muscadelle, Petit Manseng, Prunelard, Petit Verdot) had no apoplectic or dead arms. The maximum incidence, exceeding 20%, was recorded for Saperavi. The distribution of dieback phenotypes differed between cultivars. For example, Castets was characterised by a higher incidence of apoplectic arms than of dead arms, whereas the opposite pattern was observed for Chasselas (Figure 3C). Berry colour had no significant effect on the incidence of either esca foliar symptoms or plant dieback (*p* = 0.95 and *p* = 1, respectively; Figure 3A and 3C).

Finally, the incidences of esca foliar symptoms and dieback were strongly correlated for each cultivar (r = 0.72; *p* = 10^−8^; Supplementary Figure S2A). This relationship was strongest in years with an incidence of dieback of more than 4% (from 2021 to 2023), whereas it was marginal or non-significant in the preceding years (Figure 4). Despite this significant correlation, we were able to identify several cultivars with a greater expression of foliar symptoms than of dieback phenotypes, including Cabernet-Sauvignon, Sauvignon Blanc, Cabernet Franc and Morrastel. Several cultivars, including Castets, Marselan and Tinto Cao, displayed the opposite behaviour (Figure 3A and C).

**Figure 4.**
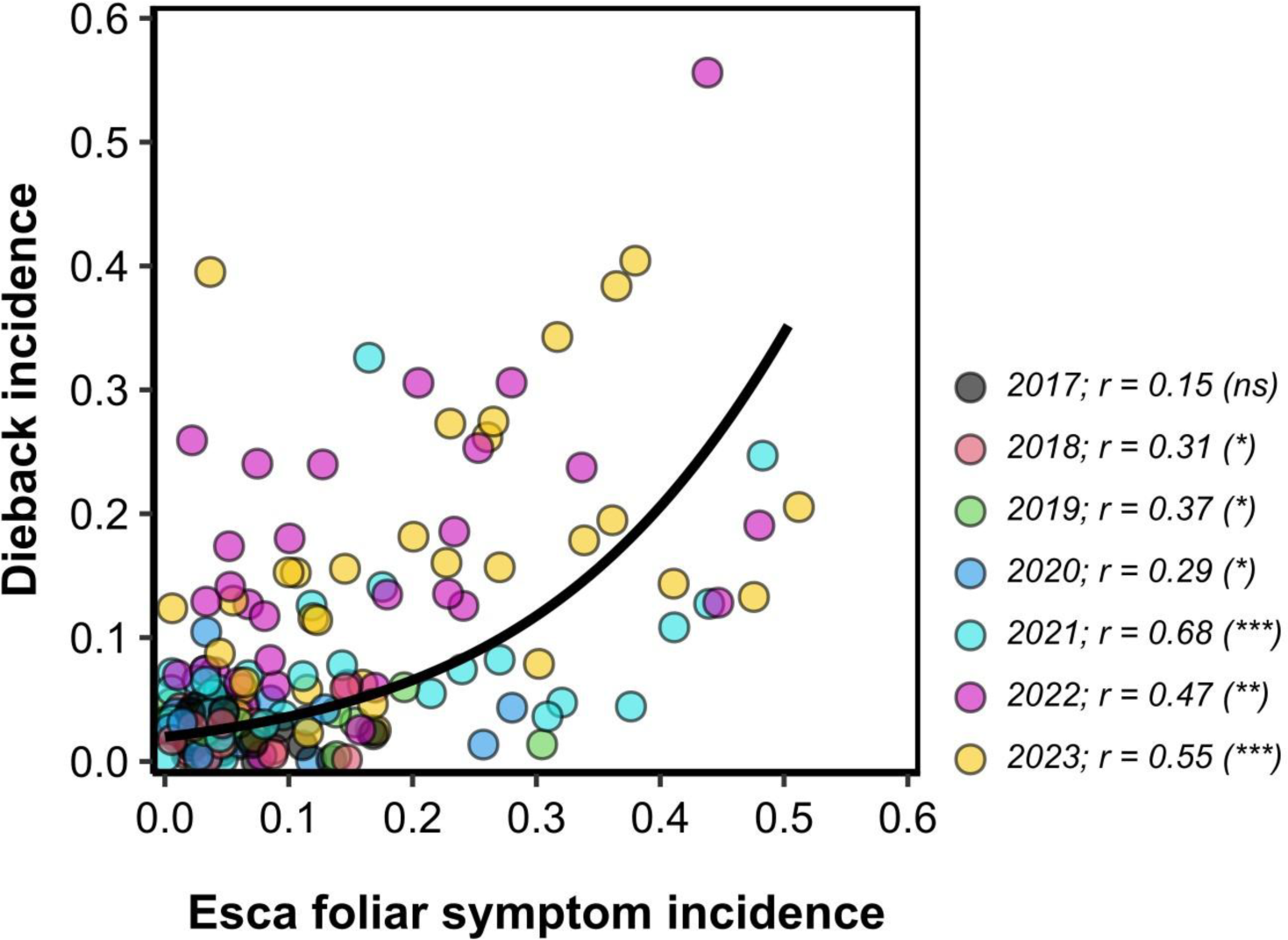
Relationship between the incidence of esca foliar symptoms and the incidence of dieback. Each dot corresponds to the value for a cultivar monitored during a single year. The regression line corresponds to a binomial model with a cloglog link fitted for all years (estimate = 6.11; *p* < 10^−16^). Dots are coloured according to the year.

By contrast, foliar symptom severity was not significantly correlated with either symptom incidence (r = 0.22, *p* = 0.14; Supplementary Figure S2B) or dieback incidence (r = 0.26, *p* = 0.08; Supplementary Figure S2C).

### The relationships between varietal patterns of susceptibility and genetic distances between cultivars are of borderline significance

Ward’s genetic distances were calculated to test for the existence of a phylogenetic signal in the epidemiological metrics. The clustering based on these distances was highly consistent with known cultivar lineages. For esca foliar symptom incidence, a significant phylogenetic signal was highlighted at global level (I = −0.10; *p* = 0.03). For this trait, phylogenetic autocorrelation values were positive for short phylogenetic distances (Figure 5A). However, this signal was not very robust, had an extremely wide confidence envelope, and was significant only for very short phylogenetic distances (Figure 5A). The distribution of the local Moran’s index I_i_ over the phylogeny revealed two hotspots of phylogenetic positive autocorrelation (i.e. nodes at which the local Moran’s index value I_i_ was significant) (Figure 5B): (i) a node of six cultivars with low incidences (Prunelard, Cot, Tannat, Petit Verdot, Petite Arvine, and Cornalin); (ii) a node of cultivars with high incidences, three of which displayed a significant signal (Sauvignon Blanc, Morrastel and Carmenere). By contrast, this method revealed no phylogenetic autocorrelation for either foliar symptom severity (I = −0.02; *p* = 0.52) or the incidence of dieback (I = −0.02; *p* = 0.30).

**Figure 5.**
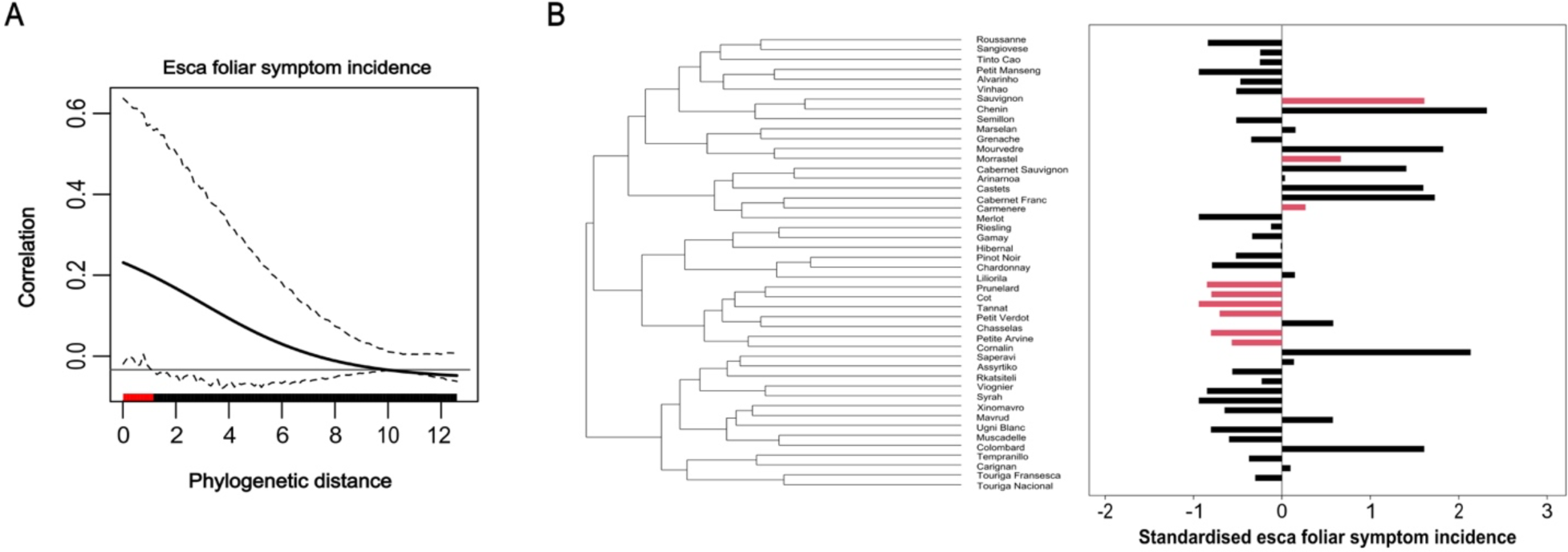
Relationship between genetic distance and the incidence of esca foliar symptoms for a panel of 46 grapevine cultivars. (A) Phylocorrelogram showing the change in Moran’s I index (measuring phylogenetic autocorrelation) along a gradient of phylogenetic distances. Black lines represent Moran’s I value. Dashed lines correspond to a confidence envelope at 95%, estimated from 1,000 bootstrap replicates. (B) Detection of the phylogenetic signal at a local scale over the phylogeny, by LIPA analysis. The dendrogram displaying genetic distances between cultivars is based on a set of 20 SSR markers. Bars correspond to standardised values (i.e. centred and scaled) for the mean incidence of esca foliar symptoms. Red bars correspond to tips for which the local Moran’s index I_i_ was significant (i.e. *p* < 0.05 in a test based on permutations).

### Varietal patterns of esca incidence remain constant over time

We investigated whether the incidence and severity of esca and plant dieback remained constant over time by performing two-by-two correlation analyses at cultivar level between the three epidemiological metrics, using the mean values for each of the six years of monitoring (a positive correlation indicates similar varietal patterns of incidence). For the incidence of foliar symptoms, all but one of the year-by-year correlations were clearly positive and significant, the exception being 2017 vs. 2022 (Figure 6A). For dieback incidence, all correlations were positive and all but five were significant (Figure 6C). By contrast, no clear inter-annual correlation pattern was identified for foliar symptom severity. Only three pairs of years displayed significant correlations, negative in one case (2018 vs. 2019) and positive in the other two (2017 vs. 2022 and 2021 vs. 2023) (Figure 6B).

**Figure 6.**
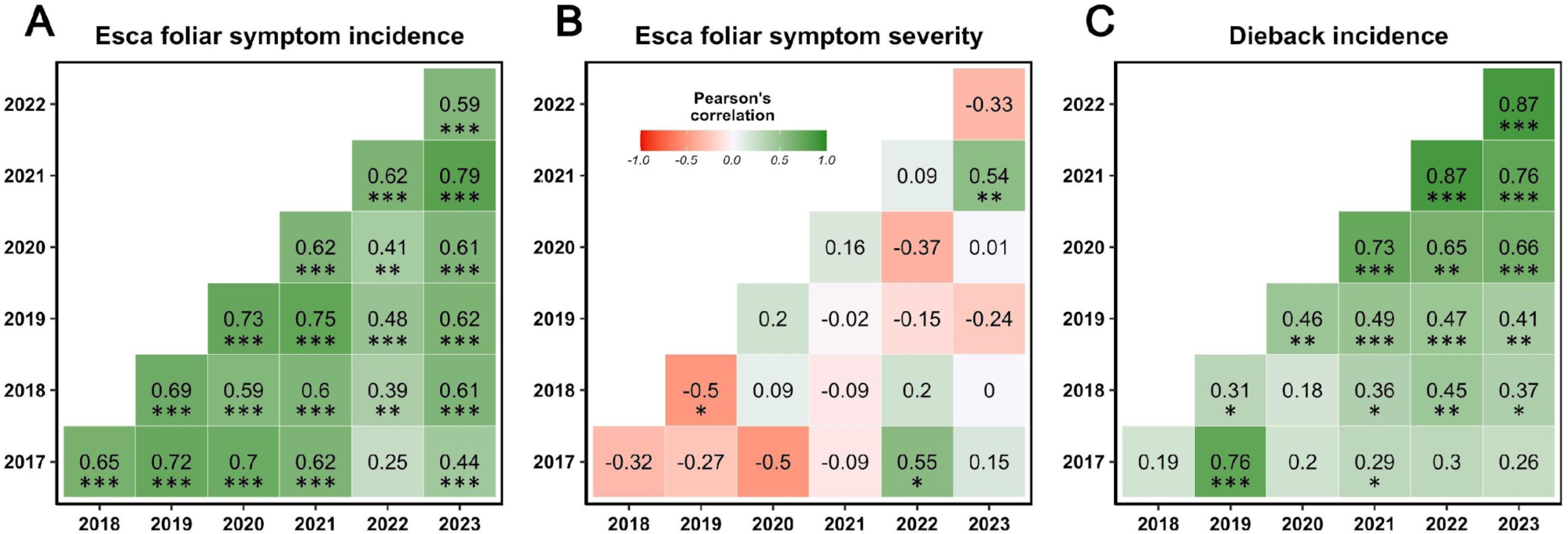
Inter-annual correlations of mean varietal values for the three variables related to esca foliar symptoms and dieback. (A) Foliar symptom incidence; (B) Severity of esca foliar symptoms, i.e. mean severity index (according to the rating scale described in Supplementary Material 2) calculated for the set of symptomatic plants; (C) Plant dieback incidence. Pearson’s correlation coefficients marked with asterisks are significant at the 5% level; *: *p* < 0.05; **: *p* < 0.01; ***: *p* < 0.001.

### A low incidence of esca is associated with high water use efficiency and low vigour at cultivar scale

The relationships between the three epidemiological metrics and cultivar characteristics were explored by measuring six phenological and ecophysiological traits in the same vineyard: bud burst, flowering and veraison dates, δ^13^C, N-tester values at flowering, and pruning weight. The variability of these traits is presented in Supplementary Table S4. Globally, the range of variability for these traits was narrow, with RSD values ranging from 3.6% (for veraison date) to 30.2% (for pruning weight). Flowering date was positively correlated with both bud burst and veraison dates (r = 0.6 and r = 0.5, respectively; Supplementary Figure S3); we therefore retained flowering date as the only phenological variable.

Correlations between the mean values of the three epidemiological metrics and these four other phenotypic traits were tested at cultivar level for the period 2017-2022 (Figure 7). δ^13^C value at harvest was significantly negatively correlated with foliar symptom expression, albeit weakly (r = 0.33; *p* = 0.03; Figure 7A). Similarly, this trait was significantly negatively correlated with dieback incidence (r = −0.32; *p* = 0.03; Figure 7I). In other words, cultivars with high water-use efficiency (i.e. a less negative δ^13^C) were less susceptible to the expression of esca foliar symptoms and dieback phenotypes. However, when this relationship was considered for each year separately, the correlation between δ^13^C and the incidence of foliar symptoms was significant only for 2022 (Supplementary Figure S4).

**Figure 7.**
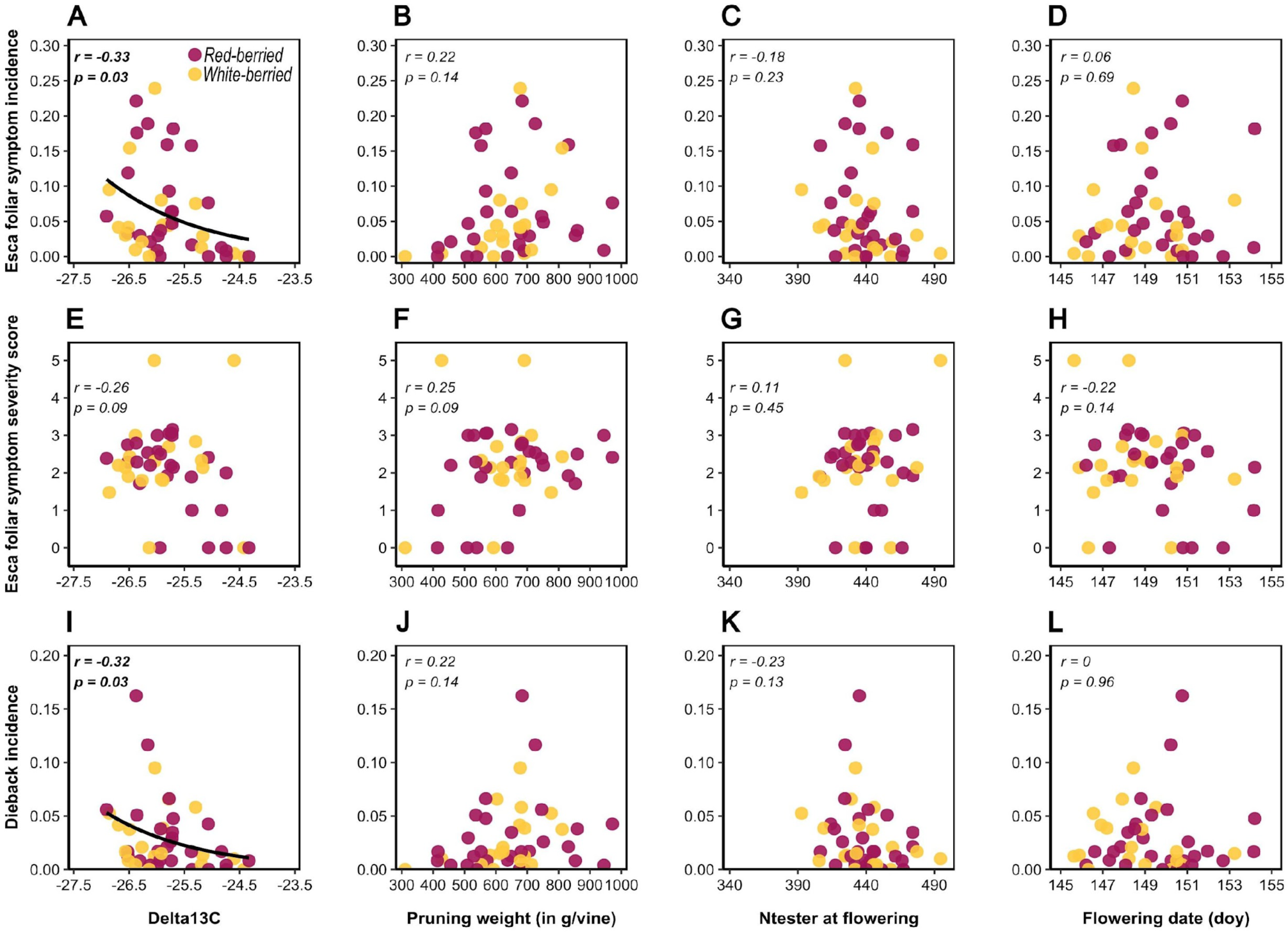
Relationships between three variables related to esca foliar symptom and dieback and four ecophysiological and phenological traits. Each dot corresponds to the value for a specific cultivar averaged over all years and blocks for the period 2017-2022. Pearson’s correlation coefficients are indicated for each relationship. The regression line is provided for significant correlations only and corresponds to a binomial model with a cloglog link as follows: (A) estimate = −0.61; *p* < 10^−16^; (E) estimate = −0.63; *p* = 10^−11^. White-berried cultivars are shown in light yellow and red-berried cultivars are shown in dark purple.

The relationship between the incidence of foliar symptoms and cultivar pruning weight was positive, but not significant (r = 0.22; *p* = 0.14; Figure 7B). Despite the weak nature of this relationship, none of the least vigorous cultivars had a high incidence of foliar symptoms.

There was a moderate negative correlation between these two ecophysiological traits (δ^13^C and pruning weight) at cultivar level (r = −0.49; *p* = 10^−5^; Supplementary Figure S3).

N-tester values at flowering and cultivar phenology (flowering date) were not significantly correlated with any of the three epidemiological metrics (Figure 7). All 12 correlations presented in Figure 7 were also tested at the subplot scale and similar trends were observed (*data not shown)*.

## Discussion

In this study, we used a unique common garden experimental vineyard to assess inter-variety variability in the expression of the vascular disease esca in grapevine. Our results indicate diverse patterns of disease susceptibility among cultivars, with some cultivars displaying no foliar symptoms in any of the seven years studied and others highly susceptible. We also identified key ecophysiological traits correlated with susceptibility to esca disease, such as cultivar water use efficiency (WUE) and vigour.

The results obtained with VitAdapt indicate that there is a high level of fairly stable cultivar-associated variability in terms of esca expression. The long-term monitoring of the VitAdapt common garden experimental vineyard revealed a significant variability of esca foliar symptoms and plant dieback between the 46 grapevine cultivars. Several cultivars, only very rarely expressed foliar symptoms or dieback phenotypes, whereas others, including several of high economic importance, were much more strongly affected by esca and dieback. The use of a common garden design, even with a limited number of plants per cultivar (n = 40), and the inclusion of a block effect in statistical models, made it possible to prevent biases in the calculation of esca incidence due to the variability of soils and viticulture practices. Our results provide new information for consideration in the discussions of cultivar rankings for esca susceptibility obtained in previous multiregional and multi-cultivar monitoring programmes. For example, Ugni Blanc, for which very high incidences were recorded in the Charentes region (France; Bruez *et al*., 2013), did not cluster among the most susceptible cultivars in our vineyard, in which all cultivars were subjected to the same environment and the same viticulture practices (e.g. double guyot pruning system). Bruez *et al*. (2013) showed that, for a given cultivar, a large proportion of the residual variability is due to the growing area. There are at least two possible explanations for this. First, there can be considerable climatic variation between regions. Second, the incidence of esca disease may be affected by differences in viticulture systems and practices from between regions (Lecomte *et al*., 2018). This may be the case for Ugni Blanc, which was analysed here in growth conditions different from those applying to productive contexts in the Charentes region (France), where it is cultivated for the production of Cognac.

Our findings clearly confirmed a number of well-known trends, such as the large differences between elite international cultivars, with Merlot (no foliar symptoms at all from eight to 14 years of age), Syrah and Pinot Noir displaying low levels of susceptibility to esca, whereas Cabernet-Sauvignon, Sauvignon Blanc, and Tempranillo were highly susceptible (consistent with the findings of Martínez-Diz *et al*., 2019; Csótó *et al*., 2023). However, they also provide integrative ecophysiological and esca susceptibility data for lesser-known varieties with promising tolerance to abiotic and biotic stresses. For example, Xinomavro, which never expressed esca in our vineyard, was classified as drought-tolerant by Plantevin *et al*. (2022) and is one of the latest ripening variety in VitAdapt (i.e. it is comparable to Grenache and Carignan, *data not shown*). Conversely, Saperavi was found to be highly prone to the expression of esca and dieback (apoplexy and death), and was also classified as susceptible to drought (Plantevin *et al*., 2022; Lamarque *et al*., 2023). More detailed studies are now required to determine whether these preliminary findings are also valid for other commercial clones of these cultivars, and for combinations with different rootstocks, in contrasting productive contexts. More generally, the extent to which esca expression varies between different clones of a single cultivar remains unclear (Murolo and Romanazzi, 2014; Moret *et al*., 2019). Zooming out to consider the entire pool of *Vitis* spp. might also be promising, to move towards the integration of susceptibility to trunk diseases as a key trait while selecting and planting both rootstocks (Gramaje *et al*., 2010; Murolo and Romanazzi, 2014) and hybrid cultivars. Here, the only hybrid variety studied, Hibernal, was found to be of intermediate susceptibility. Comparing 104 interspecific hybrids of diversified pedigrees with 201 *V. vinifera* cultivars, Csótó *et al*. (2023) demonstrated that the interspecific hybrids were globally less susceptible to trunk diseases.

A global increase in esca expression was recorded over the course of the trial, for both foliar symptoms and plant dieback, probably due to the ageing of the plants (from 8 years old at the start of monitoring to 14 years old). The incidence of esca disease is known to increase when the plants are about 10 years of age, generally reaching a maximum in plants between the ages of 15 and 25 years (Kovács *et al*., 2017; Etienne *et al*., 2024). Moreover, our data confirm that the cumulative incidence of esca foliar symptoms was much higher than the mean annual incidence, implying that, for a given subplot, the vines presenting foliar symptoms of esca differ between years. These results, alongside with previous findings, clearly indicate that it is essential to perform monitoring over several years for field trials (Reis *et al*., 2019).

Even though the incidence of esca increased over time, the relative ranking of the varieties remained constant between years. As all cultivars were compared in similar environmental conditions, this suggests that each cultivar has a constitutive level of susceptibility that is little affected by local variations of esca disease pressure (which are, in turn, influenced by several traits relating to pathogens, plants and environment; Claverie *et al*., 2020). A first genetic locus associated with grapevine susceptibility to fungal trunk pathogens was recently identified in *V. vinifera* cv. ‘Gewurztraminer’, based on internal symptoms (proportions of total necrosis and white rot) rather than the monitoring of foliar symptoms and dieback (Arnold *et al*., 2023). However, anatomical and physiological traits may contribute to cultivar susceptibility. A high density of large vessels (>100 µm) has been associated with enhanced susceptibility to *P.chlamydospora* following artificial inoculation (Pouzoulet *et al*., 2020). Based on the results of studies on very small numbers of cultivars, we can also hypothesise that wood composition (see Rolshausen *et al*., 2008 for *Eutypa lata*, another fungal pathogen of wood) and secondary metabolism (see Lemaitre-Guillier *et al*., 2020 for *Botryosphaeria* sp. diseases) may have an effect. Another non-mutually exclusive hypothesis would be the existence of interspecific and intraspecific diversity in the microbial communities interacting with different cultivars during plant pathogenesis (Laveau *et al*., 2009; Bekris *et al*., 2021).

As the genetic proximity between plant species and populations is often associated with common susceptibilities to pathogens and pests (Gilbert *et al*., 2015), we tested the hypothesis of closely related cultivars having similar patterns of susceptibility to esca. We found a significant but weak phylogenetic signal for esca susceptibility based on the incidence of foliar symptoms. This signal was associated exclusively with local nodes of closely related cultivars and was similar for all four clustering methods tested *(data not shown)*. This signal was not as robust as that likely to arise at broader levels of grouping, and may be strongly affected by the choice of genetic markers. Based on a local-scale analysis, the main hotspots of phylogenetic signal corresponded to a group of weakly susceptible cultivars. Pedigrees were available for only two of these six cultivars, Cot being an offspring of Prunelard in the VIVC. The relationships between the remaining varieties were less clear, and they even came from contrasting locations (Bacilieri *et al*., 2013). We can hypothesise that traits reducing susceptibility to esca in grapevine cultivars are converging within this group of interest. Susceptible cultivars were overrepresented at another node of the phylogeny: Sauvignon Blanc, Chenin, Marselan, Mourvedre, Morrastel, Cabernet-Sauvignon, Castets, Cabernet Franc, Carmenere. All these cultivars originate from Western and Central Europe, with six originating from South-Western France (Bacilieri *et al*., 2013). Most belong to the Savagnin and Cabernet Franc families. Nevertheless, the signal on these branches appeared unstable (i.e. significant for only three cultivars). Moreover, phylogenetic correlation did not apply to all members of a group or lineage, as represented in the vineyard studied. Thus, despite promising findings, it is difficult to conclude that esca susceptibility is clearly driven by phylogenetic patterns. Here, the genetic panel studied consisted of cultivars belonging from a single species, *V. vinifera* L. (with the exception of the *Vitis* sp. hybrid ‘Hibernal’) and was not truly representative of the overall diversity of grapevine. The search for genetic markers of trunk disease incidence would probably benefit from studies of specifically designed diversity panels (e.g. Nicolas *et al*., 2016).

Interestingly, the incidences of foliar symptoms and dieback were strongly and consistently correlated between cultivars and years. This suggests that the cultivars with a high incidence of esca were also those with a high proportion of apoplectic or dead plants. This finding echoes the work of Guérin-Dubrana *et al*. (2013) in the Bordeaux region, which showed that, in Cabernet-Sauvignon, mortality was higher in plants that had expressed esca symptoms in previous years. Nevertheless, other studies have shown that this correlation is not always valid at plant level in individual cultivars. For instance, Andreini *et al*. (2014) noted a discordance between esca symptom expression and plant mortality, especially for Cabernet-Sauvignon. Dewasme *et al*. (2022) suggested that the impact of esca on the mortality of this cultivar might be overestimated. Investigations of the mechanisms underlying foliar symptoms and plant decline, especially at vascular level, would be a promising approach to clarifying this issue.

Conversely, a lack of correlation was noted between the metrics reflecting disease incidence and those reflecting the severity of foliar symptoms. There was also no clear varietal or temporal pattern for foliar symptom severity. We hypothesise that a number of different mechanisms and factors underlie these differences in severity levels. The link between incidence and severity has long been called into question in plant science. These two traits are known to be positively correlated for a large number of airborne fungal diseases (Seem, 1984). Nevertheless, no such relationship would be expected for wilt diseases with symptoms occurring in organs (here, in leaves) distal to the infection area (here, woody organs; Seem, 1984). Few studies have specifically tested this hypothesis for trunk disease pathosystems, although several previous studies concluded that there was a positive correlation between incidence and severity in a single (Calzarano *et al*., 2018) or multiple cultivars (Romanazzi *et al*., 2009).

There have been many studies of the impact of esca disease on grapevine physiology, including plant growth (Gramaje *et al*., 2010; Dell’Acqua *et al*., 2023), photosynthesis, water relations (Magnin-Robert *et al*., 2011; Bortolami *et al*., 2021a,b), and phenology (Andreini *et al*., 2013), but very little effort has, as yet, been devoted to the opposite issue: the ecophysiological traits predicting esca incidence or severity. Here, we found that a low incidence of esca appeared to be associated with high water use efficiency and low vigour at the cultivar scale. At this same scale, δ^13^C levels were significantly negatively related to the incidences of both foliar symptoms and dieback. This indicator, averaged over several years of contrasting water availability, is used as a proxy for varietal WUE (Bchir *et al*., 2016). Cultivars with higher WUE, likely associated with lower levels of stomatal opening (Bota *et al*., 2016), were found to be less prone to the expression of esca symptoms. The underlying mechanism may be at least partly related to that proposed by Bortolami *et al*. (2021b) to elucidate the antagonistic effects of drought and esca. Genotypes with a high WUE control their transpiration much more strongly, thereby slowing the translocation of toxins from the trunk to the leaves via the vascular apparatus. Di Marco and Osti (2008) drew similar conclusions for wood decay in kiwifruit (*Actinidia deliciosa* var. deliciosa). However, in our case, this relationship was found to be unstable over time. The strongest correlation was observed in 2022, a year combining high esca intensity and low water availability. Indeed, it is known that the differences in WUE between varieties can be exacerbated by drought conditions (Plantevin *et al*., 2022), particularly when different plant genotypes are planted in the same vineyard.

Water availability is considered a driver of esca disease expression. High water levels are considered to activate esca expression (Bortolami *et al*., 2021b; Monod *et al*., 2023). In this study, the overall increase in esca incidence over time (with the ageing of the vines) made it difficult to decipher precisely the role of the water conditions in each year. Our findings suggest that, in our experimental context, plant age makes a much greater contribution to esca incidence than plant water use. Etienne *et al*. (2024) demonstrated that esca incidence reaches its maximum between 15 and 39 years old, for most of the cultivars. Several factors related either to fungal colonisation (e.g. number of infection cycles, wood microbiome) or to plant defence response (e.g. metabolism, wood properties) may potentially explain these differences related to plant age (Dissanayake *et al*., 2018; Fischer and Peighami-Ashnaei, 2019).

The least vigorous cultivars appear to be grouped among the cultivars least susceptible to esca. This relationship has already been suggested for trunk diseases, but never clearly demonstrated. It may reflect the narrower xylem vessels of less vigorous vines, increasing compartmentalisation efficiency and decreasing susceptibility to wood pathogens (Pouzoulet *et al*., 2020). We found that δ^13^C levels and pruning weight were negatively correlated. In particular, a set of weakly susceptible cultivars of low vigour were found to have less negative values for δ^13^C (e.g. Petit Manseng, Tannat, Mavrud, Colombard). We suggest that the tandem action of these two traits in the prediction of esca incidence is not a coincidence. As discussed in previous studies, these cultivars are more able to decrease their hydraulic conductivity markedly and rapidly in response to non-optimal water conditions, thus ensuring larger hydraulic safety margins but constraining plant growth and productivity (Torrez-Ruiz *et al*., 2024). Nevertheless, this relationship is ambiguous, and depends on water availability. Our findings are consistent with those of Tambussi *et al*. (2007), who found that, for common cereals, plants with a lower WUE have a competitive advantage for growth and productivity in climates with rainfall occurring during the growing season (here, a temperate oceanic climate).

Esca disease expression is also considered to be positively correlated with leaf nitrogen status (Calzarano *et al*., 2009; Li, 2015) through a roughly twofold mechanism that has already been discussed for plants (e.g. Mur *et al*., 2017; Sun *et al*., 2020) involving modifications to plant metabolism (e.g. enhanced vigour) and a favouring of fungal development and pathogenicity. In our experimental vineyard, normalised nitrogen index was not a determinant of varietal susceptibility to esca. As nitrogen status and vigour were only weakly correlated in this dataset (Supplementary Material 7), it seems likely that the variability of nitrogen status was limited in this trial, potentially accounting for the lack of correlation between vine nitrogen status and esca symptoms. This result and those of Monod *et al*. (2023), who demonstrated a negative correlation between nitrogen status and esca severity in a single cultivar growing in contrasting conditions, claim for more research, across a wide range of genotypes and growing conditions, to explore the link between vine mineral status and the incidence of trunk diseases.

Finally, we found no significant effect of phenology on the variability of esca expression between varieties. There have been few studies of the links between trunk diseases and phenology. Serra *et al*. (2018) assumed that phenology affects esca expression in relation to climatic variables. As foliar symptoms emerge only between fruit set and veraison and accumulate through the summer season in a systematic progressive sigmoidal pattern (Lecomte *et al*., 2024), we can hypothesise that varietal shifts in phenological stage may influence cultivar-specific patterns of esca expression, particularly when climatic conditions fluctuate sharply. Monitoring over a time scale finer than one year would improve our understanding of the interaction between grapevine phenology and disease dynamics.

## Conclusion

This study, making use of a unique experimental facility, adds to our knowledge of the contribution of genetic diversity to the sustainable management of grapevine dieback. Our findings confirm the utility of performing pluriannual monitoring in natural conditions, within common garden experimental vineyards, for accurate comparisons of susceptibility to trunk diseases between cultivars. The ranking of varieties for susceptibility, based on the incidence of esca foliar symptoms, was consistent with the findings of a previous field trial conducted in Italy (Murolo and Romanazzi, 2014; Figure 8A). A weaker correlation was also identified with the ranking obtained in a network of French vineyards (Figure 8B). Correlations were tested with the ranking of varietal susceptibility to other grapevine diseases. The susceptibility to *flavescence dorée*, caused by a phytoplasma, was significantly correlated with esca susceptibility (eight cultivars in common with VitAdapt; τ = 0.76; p = 10^−3^; Eveillard *et al*., 2016), while the susceptibility to the fungal trunk disease Eutypa dieback was not (29 cultivars in common; τ = 0.23; p = 0.11; Sosnowski *et al*., 2022). No correlation was found on VitAdapt between the incidence of esca foliar symptoms and the incidence of symptoms associated with *Botrytis cinerea* (an aerial necrotrophic fungus) on bunches at harvest (22 cultivars in common; τ = −0.22; p = 0.15; Paňitrur-De La Fuente *et al*., 2018). Interestingly, no correlation was found between the ranking of varieties for the incidence of esca foliar symptoms in VitAdapt and the susceptibility gradients obtained by artificially inoculating cuttings with wood pathogens such as *P. chlamydospora* (Pouzoulet *et al*., 2020; Figure 8C) or *Neofusicoccum parvum* (*unpublished data*; Figure 8D). This confirms that the susceptibility of grapevine genotypes to trunk diseases cannot be assessed solely on the basis of inoculations under control conditions, as already discussed by several authors (e.g. Sosnowski *et al*., 2007; Reis *et al*., 2019).

**Figure 8.**
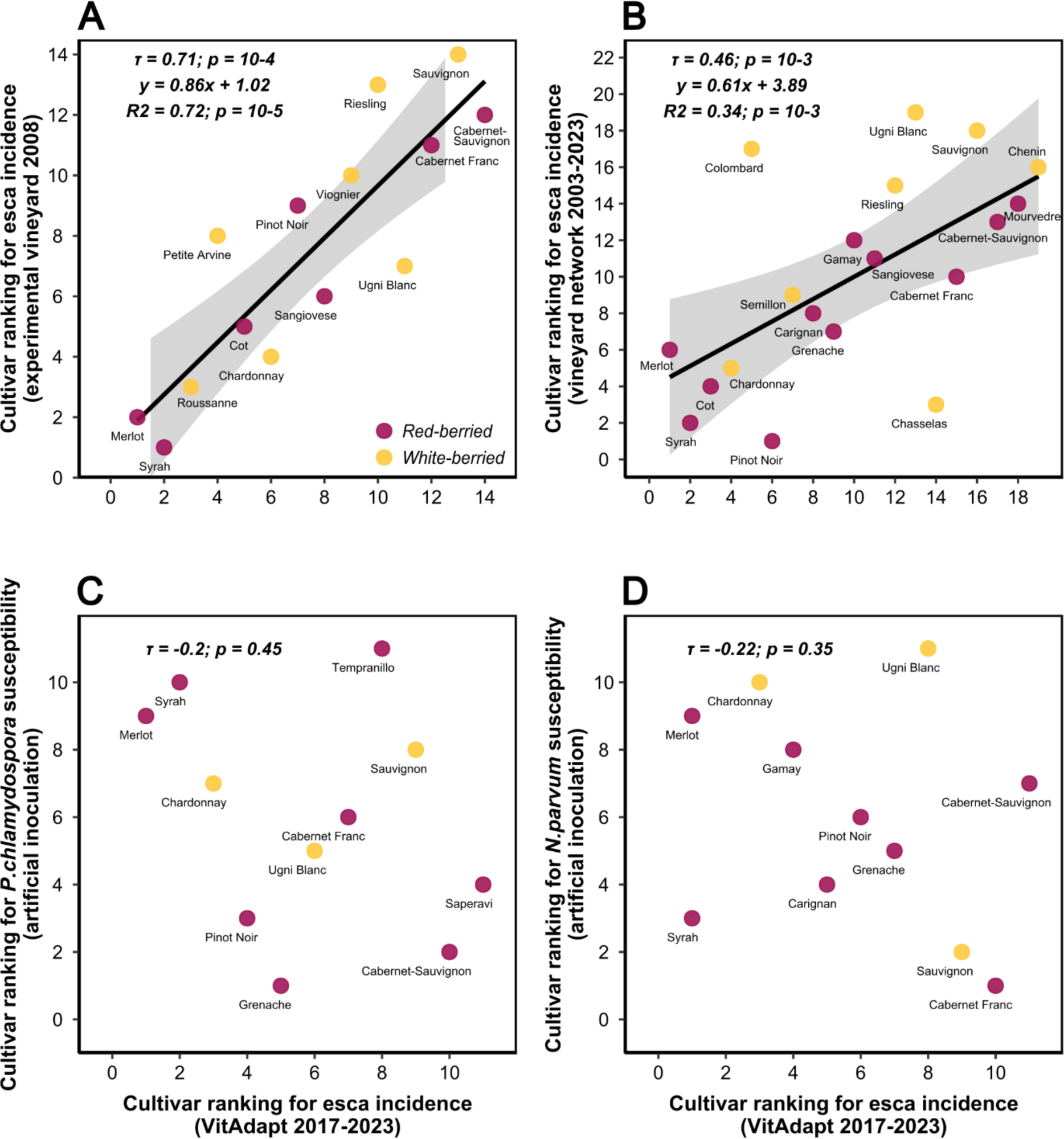
Scatterplots showing the relationships between cultivar ranks for susceptibility to esca and wood pathogens as assessed in this work and reported in previous publications. Each dot corresponds to the ranking of a cultivar in a given study, for the range of cultivars common to the two studies compared. (A) Relationship between the cultivar rankings obtained in this study and those obtained by Murolo and Romanazzi (2014) in a one-year (2008) study monitoring of esca foliar symptom expression in an experimental vineyard in Italy (14 cultivars common to the two studies). The regression line corresponds to a linear model, and the standard error is provided; (B) Relationship between the cultivar rankings obtained in this study and those obtained by Etienne *et al*. (2024), in a study over six years (2003-2023) monitoring esca foliar symptom expression in a network of 2084 vineyards located in French grape-producing areas (19 cultivars common to the two studies); (C) Relationship between the cultivar rankings obtained in this study and those obtained by Pouzoulet *et al*. (2020) by phenotyping the length of stem necrosis induced by *Phaeomoniella chlamydospora* after 12 weeks of incubation (11 cultivars common to the two studies). Rank-based Kendall’s correlation coefficients are indicated for each relationship; (D) Relationship between the cultivar rankings obtained in this study and those obtained by phenotyping the length of stem necrosis induced by *Neofusicoccum parvum* after 28 weeks of incubation (*unpublished data;* 11 cultivars common to the two studies). Rank-based Kendall’s correlation coefficients are indicated for each relationship.

Furthermore, susceptibility to trunk diseases is potentially plastic in response to the environment. To facilitate the selection of the most suitable cultivars for each production context, it therefore seems necessary to combine (i) replication of such monitoring under contrasting conditions and (ii) modelling-based approaches, integrating epidemiological, physiological and pedoclimatic variables. Within this framework, a complementary strategy would involve monitoring a single cultivar in multiple environments (see Monod *et al*., 2023). Finally, we recommend tackling the highly multifactorial nature of plant dieback through integrative approaches taking multiple factors, such as other pests and diseases, climatic hazards, soil fertility and technical management, into account.

## Acknowledgements

We would like to thank the Experimental Viticultural Unit of Bordeaux 1442, INRAE, F-33883 Villenave d’Ornon for managing the experimental vineyards and collecting data. We thank the teams from SAVE and EGFV for data collection, Gwenaëlle Comont (SAVE) and Dr Mark Sosnowski (SARDI) for providing data used to correlate varietal susceptibility rankings (*N. parvum* inoculations and Eutypa dieback, respectively). We also thank Julie Sappa for proofreading and English editing. Pierre Gastou’s PhD grant was awarded by the French *Ministère de l’Enseignement Supérieur et de la Recherche*. This study received financial support from Château-Figeac (Saint-Emilion), the French government in the framework of the IdEX Bordeaux University “Investments for the Future” programme / GPR Bordeaux Plant Sciences), and the French National Research Agency (ANR) in the framework of the “Investments for the Future Programme”, within the Cluster of Excellence COTE (ANR-10-LABX-45). The long-term monitoring of the Vitadapt vineyard was supported by INRAE, *Région Nouvelle-Aquitaine*, the VITGREFSEC (22003182) and VIFADEPT (22001762) projects (*Conseil Interprofessionnel des Vins de Bordeaux* - CIVB), the projects PHYSIOPATH (22001150) and ESCAPADE (22001436) (“*Plan National Dépérissement du Vignoble*” programme, FranceAgriMer/CNIV) and was conducted as part of the International Associated Laboratory (LIA) Innogrape project.

