## Supplementary materials for "Large gradient of susceptibility to esca disease revealed by long-term monitoring of 46 grapevine cultivars in a common garden vineyard"

**SUPPLEMENTARY TABLE S1. List of plant material included in this study: 45 *Vitis vinifera* L. cultivars and a hybrid (Hibernal)**

General descriptive information for each of the cultivars is given in the Vitis International Variety Catalogue (VIVC, www.vivc.de) for berry colour and country of origin; and by Bacilieri *et al.* (2013) for genetic prole. The origin and plantation date of the plant material was provided by Destrac Irvine and van Leeuwen (2016).

The origin of the plant materials was as follows: INRA Bx = Domaine de la Grande Ferrade, Centre INRAE Nouvelle Aquitaine Bordeaux, 33882 Villenave d'Ornon, France; ENTAV = Etablissement National Technique Amélioration Viticulture, 30240 Le Grau-du-Roi, France; Vassal = Domaine de Vassal INRA, 34340 Marseillan, France; CA 33 = Chambre d’Agriculture Gironde - Vinopôle Bordeaux Aquitaine, 33295 Blanquefort, France; CA 64 = Chambre d’Agriculture Pyrénées-Atlantiques, 64000 Pau, France; CA 11 = Chambre d’Agriculture Aude, 11570 Palaja, France; Geisenheim = Hochschule Geisenheim University, 65366 Geisenheim, Germany; Gaillac = Institut Français de la Vigne et du Vin, V'Innopôle Sud-Ouest, 81 310 Peyrole, France).

| **Cultivar name** | **Berry colour** | **Country of origin** | **Genetic prole** | **Plant material**  **origin** | **Plantation date** |
| --- | --- | --- | --- | --- | --- |
| Alvarinho | White | Portugal | occidentalis | *NA* | 2010 |
| Arinarnoa | Red | France | occidentalis | INRA BX | 2009 |
| Assyrtiko | White | Greece | *NA* | Vassal | 2009 |
| Cabernet-Sauvignon | Red | France | occidentalis | CA 33 | 2009 |
| Cabernet Franc | Red | France | occidentalis | CA 33 | 2009 |
| Carignan | Red | France | occidentalis | ENTAV | 2009 |
| Carmenere | Red | France | occidentalis | CA 33 | 2009 |
| Castets | Red | France | occidentalis | INRA BX | 2009 |
| Chardonnay | White | France | occidentalis | INRA BX | 2009 |
| Chasselas | White | France | orientalis | INRA BX | 2009 |
| Chenin | White | France | occidentalis | ENTAV | 2009 |
| Colombard | White | France | occidentalis | CA 33 | 2010 |
| Cornalin | Red | Switzerland | occidentalis | INRA BX | 2009 |
| Cot | Red | France | occidentalis | CA 33 | 2009 |
| Gamay | Red | France | occidentalis | ENTAV | 2009 |
| Grenache | Red | Spain | orientalis | ENTAV | 2009 |
| Hibernal | White | Germany | *NA* | Geisenheim | 2009 |
| Liliorila | White | France | occidentalis | INRA BX | 2009 |
| Marselan | Red | France | occidentalis | ENTAV | 2009 |
| Mavrud | Red | Bulgaria | pontica | *NA* | 2010 |
| Merlot | Red | France | occidentalis | CA 33 | 2009 |
| Morrastel | Red | Spain | occidentalis | ENTAV | 2009 |
| Mourvedre | Red | Spain | occidentalis | ENTAV | 2009 |
| Muscadelle | White | France | occidentalis | CA 33 | 2009 |
| Petit Manseng | White | France | occidentalis | CA 64 | 2010 |
| Petit Verdot | Red | France | occidentalis | CA 33 | 2009 |
| Petite Arvine | White | Switzerland | occidentalis | INRA BX | 2009 |
| Pinot Noir | Red | France | occidentalis | ENTAV | 2009 |
| Prunelard | Red | France | orientalis | IFV Gaillac | 2009 |
| Riesling | White | Germany | occidentalis | INRA BX | 2009 |
| Rkatsiteli | White | Georgia | orientalis | INRA BX | 2009 |
| Roussanne | White | France | occidentalis | ENTAV | 2009 |
| Sangiovese | Red | Italy | occidentalis | ENTAV | 2009 |
| Saperavi | Red | Georgia | orientalis | INRA BX | 2009 |
| Sauvignon | White | France | occidentalis | CA 33 | 2009 |
| Semillon | White | France | occidentalis | CA 33 | 2009 |
| Syrah | Red | France | occidentalis | CA 11 | 2010 |
| Tannat | Red | France | orientalis | CA 64 | 2010 |
| Tempranillo | Red | Spain | orientalis | ENTAV | 2009 |
| Tinto Cao | Red | Portugal | occidentalis | Vassal | 2009 |
| Touriga Francesa | Red | Portugal | occidentalis | Vassal | 2009 |
| Touriga Nacional | Red | Portugal | occidentalis | INRA BX | 2009 |
| Ugni Blanc | White | Italy | pontica | CA 33 | 2009 |
| Vinhao | Red | Portugal | occidentalis | *NA* | 2010 |
| Viognier | White | France | occidentalis | ENTAV | 2009 |
| Xinomavro | Red | Greece | *NA* | *NA* | 2010 |

**SUPPLEMENTARY TABLE S2. Description of the rating scale used to assess esca foliar symptoms and plant dieback**

For each code, a short description of the corresponding phenotype is supplied. The scoring system was adapted from Lecomte *et al.* 2012 (Plant Disease, 96(7), 924-934). The scores from both symptomatic arms can be averaged to describe esca foliar symptom severity at the plant level.

| **Rating** | **Description** |
| --- | --- |
| 0 | No esca symptom or plant dieback |
| 1 | Slight esca symptoms on one arm (some drying or discoloration restricted to a few leaves) |
| 3 | Pronounced esca symptoms on one arm (pronounced drying or discoloration on several stems) |
| 5 | Very severe esca symptoms on one arm (drying, discoloration and bunch wilt) |
| 7 | Apoplexy on one arm (generalised wilting) |
| 9 | Dead arm |
| 99 | Dead vine (both arms) |

**SUPPLEMENTARY TABLE S3. Description of the method used to correct N-tester values for interspecific comparisons**

Mean N-tester values at mid-flowering, across several years, are given for 45 cultivars. For each cultivar, the difference with respect to the reference cultivar (Sauvignon Blanc) is shown. Different colours indicate different groups according to an ascendant hierarchical classification (AHC). The mean correction applied to the whole set of cultivars in each group is provided.

| **Cultivar** | **Years** | **Average N-tester score at flowering** | **Difference relative to reference variety** | **Correction for the group** |
| --- | --- | --- | --- | --- |
| Chasselas | 2015-2021 | 324.2917 | +110 | **+70** |
| Grenache | 2015-2021 | 346.1591 | +88 |  |
| Xinomavro | 2020-2021 | 347.5 | +86 |  |
| Tinto Cao | 2020-2021 | 371.5 | +62 |  |
| Petite Arvine | 2020-2021 | 375.5 | +58 |  |
| Vinhao | 2015-2021 | 375.8864 | +58 |  |
| Semillon | 2015-2021 | 385.2273 | +49 |  |
| Colombard | 2015-2021 | 391.2955 | +43 |  |
| Prunelard | 2020-2021 | 397.25 | +37 |  |
| Viognier | 2015-2021 | 402.4583 | +32 | **0** |
| Carmenere | 2015-2021 | 403.5 | +30 |  |
| Marselan | 2015-2021 | 404.2273 | +30 |  |
| Touriga Nacional | 2015-2021 | 406.087 | +28 |  |
| Hibernal | 2020-2021 | 409 | +25 |  |
| Syrah | 2015-2021 | 410.1591 | +24 |  |
| Alvarinho | 2015-2021 | 411.4091 | +23 |  |
| Ugni Blanc | 2015-2021 | 416.8958 | +17 |  |
| Cot | 2015-2021 | 419.2273 | +15 |  |
| Chardonnay | 2015-2021 | 419.3409 | 15 |  |
| Sangiovese | 2015-2021 | 419.7583 | +14 |  |
| Castets | 2015-2021 | 420.0455 | +14 |  |
| Muscadelle | 2015-2021 | 423.2045 | +11 |  |
| Gamay | 2015-2021 | 429.0682 | +5 |  |
| Sauvignon | 2015-2021 | 433.6304 | 0 |  |
| Saperavi | 2015-2021 | 437.9545 | -4 |  |
| Tannat | 2015-2021 | 439.0455 | -5 |  |
| Cabernet Franc | 2015-2021 | 443.0227 | -9 |  |
| Petit Manseng | 2015-2021 | 445.1136 | -11 |  |
| Cabernet-Sauvignon | 2015-2021 | 456.2955 | -22 | **-40** |
| Chenin | 2015-2021 | 458.4375 | -24 |  |
| Riesling | 2015-2021 | 460.0909 | -26 |  |
| Mourvedre | 2015-2021 | 463.3864 | -29 |  |
| Pinot Noir | 2015-2021 | 465.8409 | -32 |  |
| Carignan | 2015-2021 | 467.3333 | -33 |  |
| Roussanne | 2015-2021 | 468.2045 | -34 |  |
| Liliorila | 2020-2021 | 469 | -35 |  |
| Morrastel | 2020-2021 | 469 | -35 |  |
| Merlot | 2015-2021 | 476.8125 | -43 |  |
| Petit Verdot | 2015-2021 | 479.7727 | -46 |  |
| Tempranillo | 2015-2021 | 481.2292 | -47 |  |
| Arinarnoa | 2020-2021 | 483 | -49 |  |
| Assyrtiko | 2020-2021 | 485.75 | -52 |  |
| Mavrud | 2020-2021 | 486 | -52 |  |
| Touriga Francesa | 2015-2021 | 494.8043 | -61 |  |
| Rkatsiteli | 2015-2021 | 513.2045 | -79 |  |
| Cornalin | *NA* | *NA* | *NA* | ***NA*** |

**SUPPLEMENTARY TABLE S4: Overall statistics for ecophysiological and phenological data in the VitAdapt common garden experimental vineyard (Villenave d’Ornon, France) between 2017 and 2022**

The statistics shown are the mean, standard deviation, relative standard deviation, minimum and maximum. For each trait, values are calculated from 1,104 observations (46 cultivars; four blocks; six years). Calendar dates were used for the analysis (day number within the year).

| **Trait** | **mean** | **SD** | **RSD** | **min** | **max** |
| --- | --- | --- | --- | --- | --- |
| Flowering date | 149.3 | 5.7 | 3.8% | 136.6 | 165 .1 |
| Bud burst date | 92.8 | 7.7 | 8.3% | 65.0 | 122.1 |
| Veraison date | 213.5 | 7.7 | 3.6% | 191.6 | 235.7 |
| δ^13^C | -25.8 | 1.38 | 5.4% | -28.8 | -22.2 |
| N-tester at flowering | 436.9 | 56.8 | 13.0% | 169.0 | 436.9 |
| Pruning weight (in g) | 645.3 | 194.7 | 30.2% | 157.5 | 1395.9 |

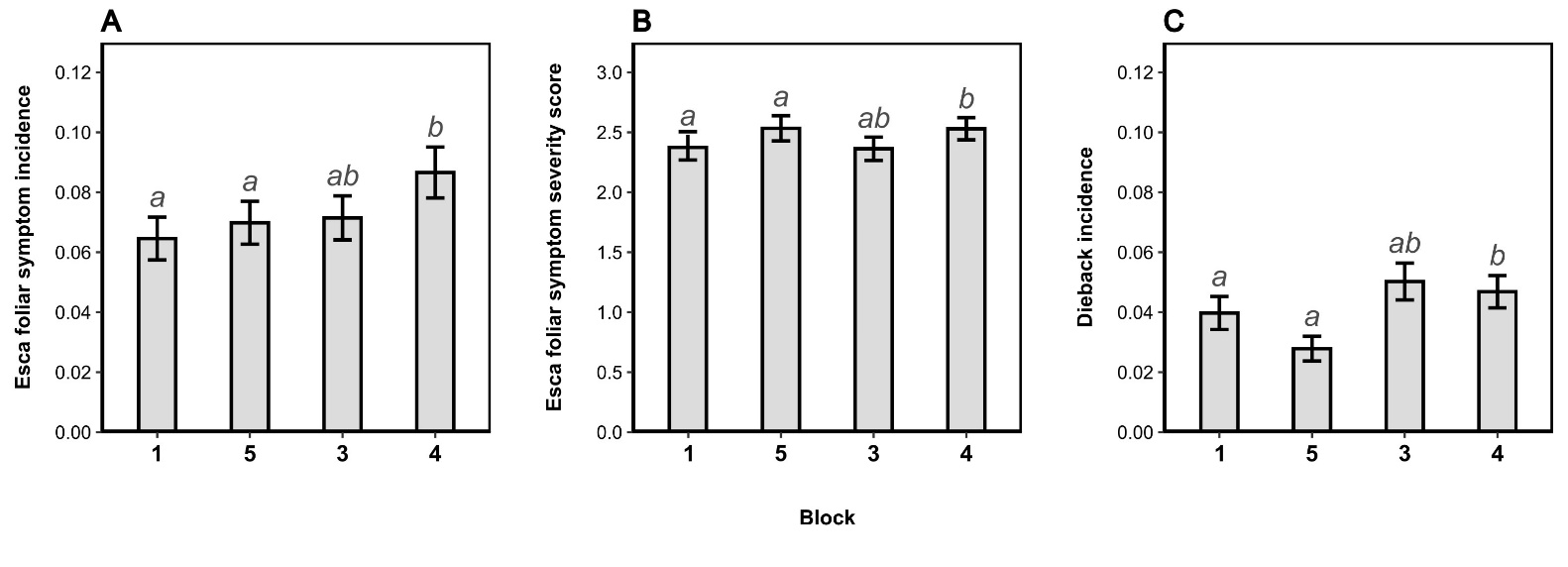

**SUPPLEMENTARY FIGURE S1. Effect of block on the esca and dieback epidemiological metrics of all cultivars for the period 2017-2023**

(A) Esca foliar symptom incidence (mean ± SEM); (B) Severity of esca foliar symptoms (mean ± SEM), i.e. mean severity index (according to the rating scale described in Supplementary Material 2) calculated for the set of plants with symptoms; (C) Plant dieback incidence (mean ± SEM). The letters correspond to the groups of significance according to Tukey’s test with an alpha risk of 5%.

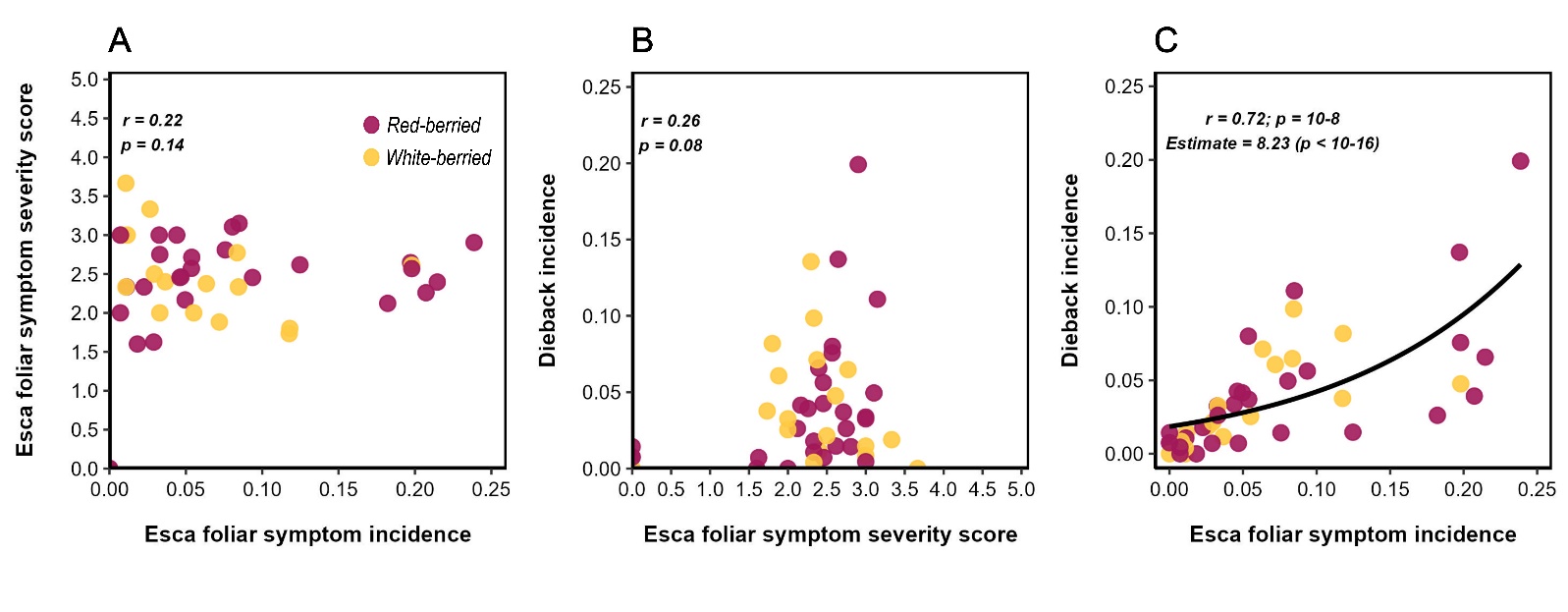

**SUPPLEMENTARY FIGURE S2. Scatterplots showing the relationships between the three esca and dieback epidemiological metrics. Each dot corresponds to the mean value for all years and blocks for a given cultivar (for the period 2017-2023)**

(A) Relationship between foliar symptom incidence and plant dieback incidence. The regression line corresponds to a binomial model with a cloglog link; (B) Relationship between foliar symptom incidence and foliar symptom severity; (C) Relationship between plant dieback incidence and foliar symptom severity. Pearson’s correlation coefficients are indicated for each relationship. White-berried cultivars are shown in light yellow and red-berried cultivars are shown in dark purple.

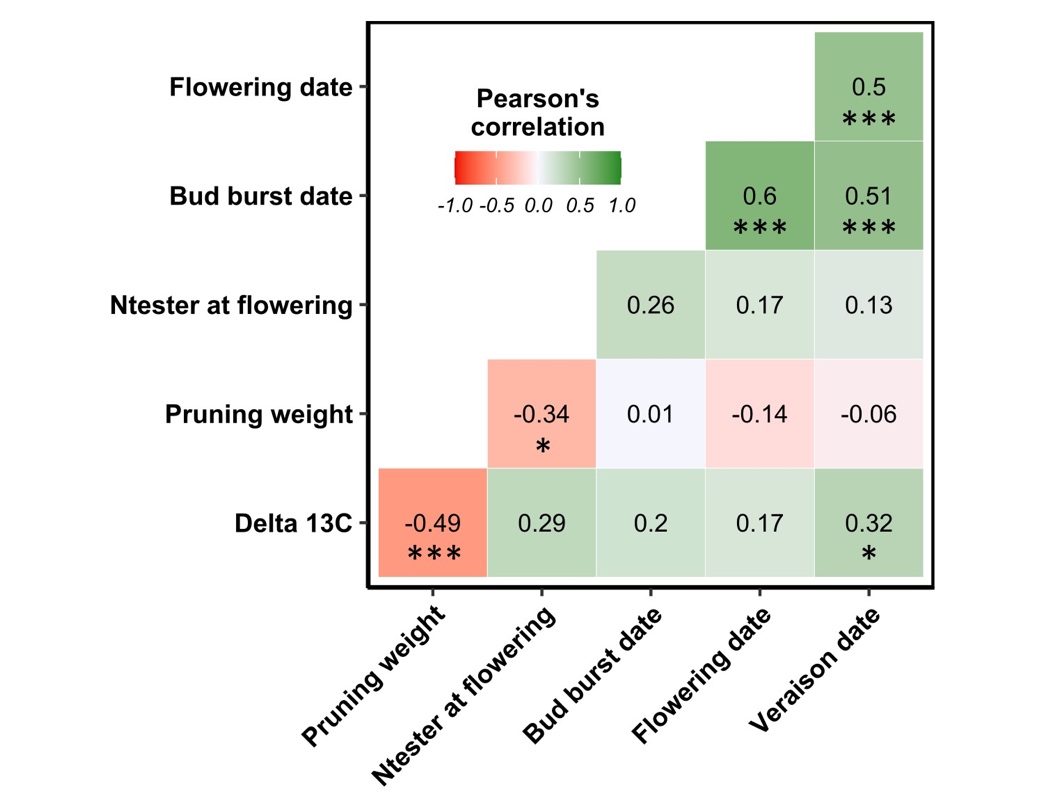

**SUPPLEMENTARY FIGURE S3. Correlations between mean varietal values for ecophysiological and phenological variables**

Pearson’s correlation coefficients marked with asterisks are significant at the 5 % threshold; *: *p* < 0.05; **: *p* < 0.01; ***: *p* < 0.001

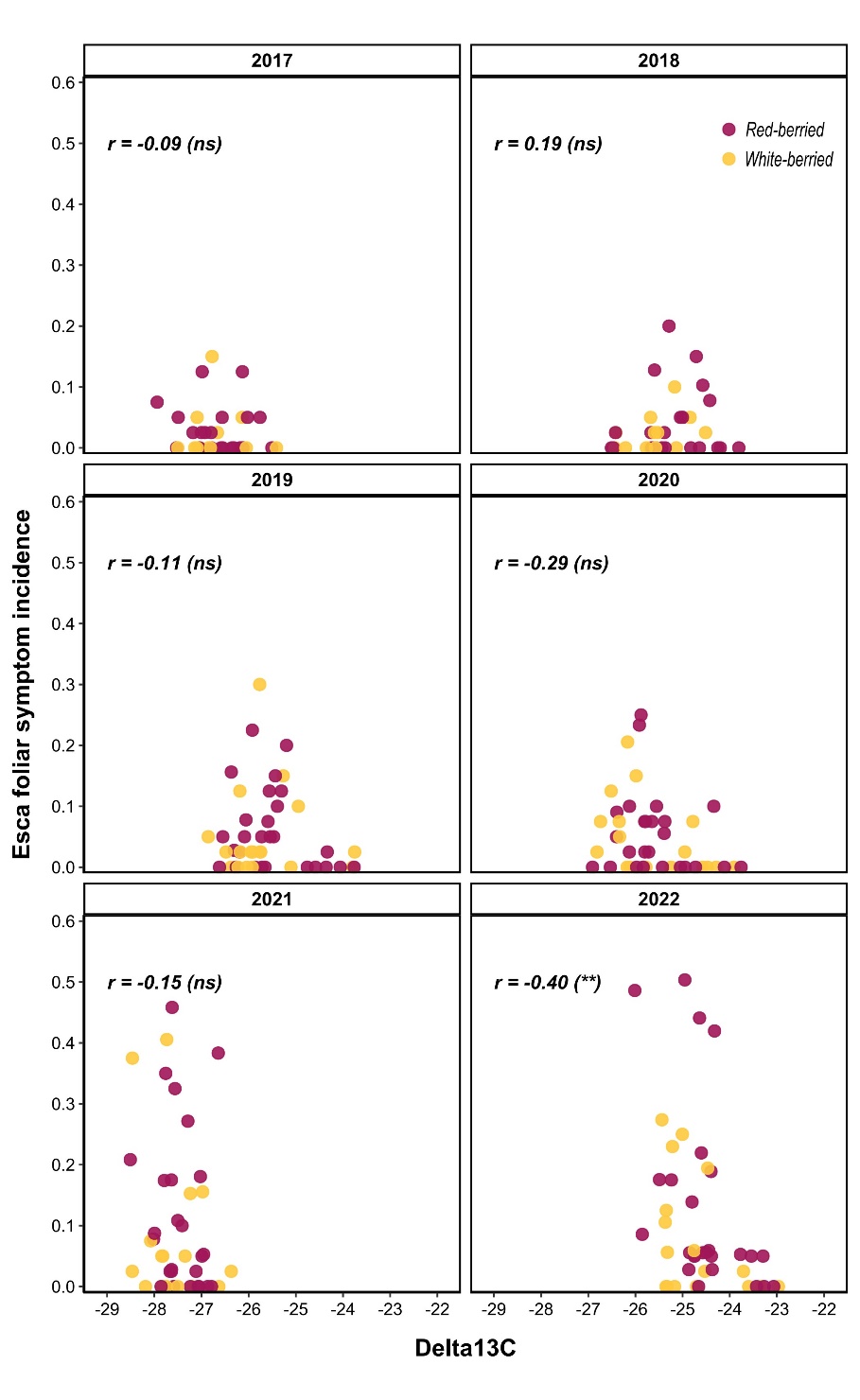

**SUPPLEMENTARY FIGURE S4. Scatterplots showing the relationships between foliar symptom incidence and δ^13^C for the various years**

Each dot corresponds to the mean value for all blocks for a given cultivar (for the 2017-2022 period). Pearson’s correlation coefficients are indicated for each relationship. White-berried cultivars are shown in light yellow and red-berried cultivars are shown in dark purple.
